# Household Members Do Not Contact Each Other at Random: Implications for Infectious Disease Modelling

**DOI:** 10.1101/220202

**Authors:** Nele Goeyvaerts, Eva Santermans, Gail Potter, Andrea Torneri, Kim Van Kerckhove, Lander Willem, Marc Aerts, Philippe Beutels, Niel Hens

## Abstract

Airborne infectious diseases such as influenza are primarily transmitted from human to human by means of social contacts and thus easily spread within households. Epidemic models, used to gain insight in infectious disease spread and control, typically rely on the assumption of random mixing within households. Until now there was no direct empirical evidence to support this assumption. Here, we present the first social contact survey specifically designed to study contact networks within households. The survey was conducted in Belgium (Flanders and Brussels) in 2010-2011. We analyzed data from 318 households totaling 1266 individuals with household sizes ranging from 2 to 7 members. Exponential-family random graph models (ERGMs) were fitted to the within-household contact networks to reveal the processes driving contact between household members, both on weekdays and weekends. The ERGMs showed a high degree of clustering and, specifically on weekdays, decreasing connectedness with increasing household size. Furthermore, we found that the odds of a contact between father and child is smaller than for any other pair except for older siblings. Epidemic simulation results suggest that within-household contact density is the main driver of differences in epidemic spread between complete and empirical-based household contact networks. The homogeneous mixing assumption may therefore be an adequate characterization of the within-household contact structure for the purpose of epidemic simulation. However, ignoring the contact density when inferring from an epidemic model will result in biased estimates of within-household transmission rates. Further research on the implementation of within-household contact networks in epidemic models is necessary.

**Significance Statement:** Households have a pivotal role in the spread of airborne infectious diseases. Households are bridging units between schools and workplaces, and social contacts within households are frequent and intimate, allowing for rapid disease spread. Infectious disease models typically assume that members of a household contact each other randomly. Until now there was no direct empirical evidence to support this assumption. In this paper, we present the first social contact survey specifically designed to study contact networks within households with young children. We investigate which factors drive contacts between household members on one particular day by means of a statistical model. Our results suggest the importance of connectedness within households over heterogeneity in number of contacts.

---

Households are crucial units in the epidemiology of airborne infectious diseases such as influenza, smallpox and SARS. Relations between household members are typically characterized by frequent and intimate contacts, allowing for rapid disease spread within the household upon introduction of an infectious case. As stated by Ferguson et al. [1]: “being a member of a household containing an influenza case is in fact the largest single risk factor for being infected oneself [2, 3]”. Furthermore, households with children have a bridging function, allowing an infection to spread from schools to workplaces and vice versa. Inference from household final-size data revealed that children play a key role in bringing influenza infection into the household and in transmitting the infection to other household members [3]. Households are the most common transmission unit used in observational studies and in epidemic models.

Many epidemic models rely on the assumption of homogeneous (random) mixing within households. In early work, Reed-Frost type of models were used to estimate household and community transmission parameters from household final size data, assuming a constant probability of infection from the community [4–6]. Ball et al. [7] generalized this to the so-called ‘households model’ with two levels of mixing, assuming random mixing within households (local) and in the entire population (global), the latter typically at a much lower rate. The analytical tractability of the households model allowed the theoretical study of epidemic phenomena. This has led to the definition of threshold parameters such as the reproduction number *R*^*^, representing the average number of households infected by a typical infected household in a totally susceptible population [7, 8]. Meyers et al. [9] used a contact network model in an urban setting incorporating households as complete networks (cliques) to explain the early epidemiology of SARS. Individual-based simulation models of infectious disease transmission incorporate detailed individual-level information in order to account for heterogeneities relevant to the application (e.g. demography, socio-economics, genetics; [10–12]). These models allow to incorporate more detailed structure in specific settings such as schools and workplaces, but typically assume random mixing in households. Studies that particularly highlight within-household transmission and control policies targeting households can be found in [13] and [1].

Until now there was no direct empirical evidence to support the assumption of homogeneous mixing within households. Egocentric contact surveys entailed partially observed within-household contact networks and only allowed for indirect inference of the unobserved network links [14, 15]. It has been argued that greater realism could be gained by considering different household compositions and contact heterogeneity within households [16].

In this paper, we describe the first social contact survey specifically designed to study contact networks within households. This study enables us to empirically assess the assumption of homogenous mixing, e.g. by studying the effect of age and gender on social distance within households. Furthermore, it provides an answer to one of the key questions from inference on household models: how does the density of the contact network scales with the household size [16]? When ignoring contact heterogeneity between household members, the contact network density equals the contact rate between two individuals in a household and is a determinant for the within-household transmission rate of airborne infectious diseases [17, 18]. Lastly, this study makes it possible to assess reporting quality for diary-reported contact surveys by looking at reciprocity, i.e. symmetry in contact reporting. We use exponential-family random graph models (ERGMs; [19]) to develop a plausible model for within-household contact networks and to gain insight in the factors driving contacts between household members. We then compare these empirically grounded ERGMs to the assumption of random mixing using stochastic simulations of an epidemic in the mise en scene of the households model with two levels of mixing.

## Results

### Household contact survey

In 2010-2011, a survey was conducted to study social contact behaviour in households with young children in Belgium (Flanders and Brussels). A larger similarly designed parallel contact survey in individuals from separate households is described elsewhere [20, 21]. Participants were recruited by random digit dialing and stratified sampling ensured representativeness in terms of geographical spread, day and week-weekend distribution, and age and gender of the youngest child. All participants were asked to anonymously complete a paper diary recording their contacts during one randomly assigned day without changing their usual behaviour

The survey focused on households with at least one child of age 12 years or less. Upon sampling, all persons living more than 50% of the time in the household were defined as household members and recruited to take part in the survey. Participants had to record all persons they made contact with during a 24 hour period assigned to them. A contact was defined as a two-way conversation at less than 3 meters distance or a physical contact involving skin-to-skin touching (either with or without conversation). The information recorded included the exact or estimated age (interval) and gender of each contacted person, physical touching (yes/no), location, frequency and total duration of the contact, and whether or not the contacted person was a household member. If they contacted someone multiple times on a day, participants specified this as a single contact, along with the estimated contact duration accumulated over the day and set the location category to ‘multiple’ if that person was contacted at two or more different locations.

From the 342 households that participated in the survey, 24 households were excluded because of missing data. We analyzed data from 318 households including 1266 participants who recorded 19685 contacts in total, with household sizes ranging from 2 to 7. Within-household contacts were identified and matched with other household members using the procedure described in the SI text. This entailed 3821 identified within-household contacts with 98% reciprocity, indicating a good quality of reporting as expected in this household setting [22]. We assumed all social contacts to be reciprocal, depicting each household as an undirected network where nodes represent household members and edges represent contacts within the household. This resulted in a total of 1946 distinct within-household contacts of which 1861 (96%) involved physical contact (Figure S1).

Figure S1 shows that contacts between household members were of long duration, which is consistent with findings from previous social contact surveys [18] and from individual-based simulation models creating so-called synthetic populations [23]. Further, interactions between household members occurred (almost) daily and 66% of household members only met each other at home on their assigned day, while 33% met at multiple locations of which 98% included home. In the following, we focus on physical contacts (with and without conversation) since it has been shown that these better explain the observed age-specific seroprevalence of airborne infections, such as varicella and parvovirus B19, compared to non-physical contacts [24–26]. Figure 1 allows to appreciate at a glance the diversity in household size and network configurations we studied through the survey.

**Fig 1.**
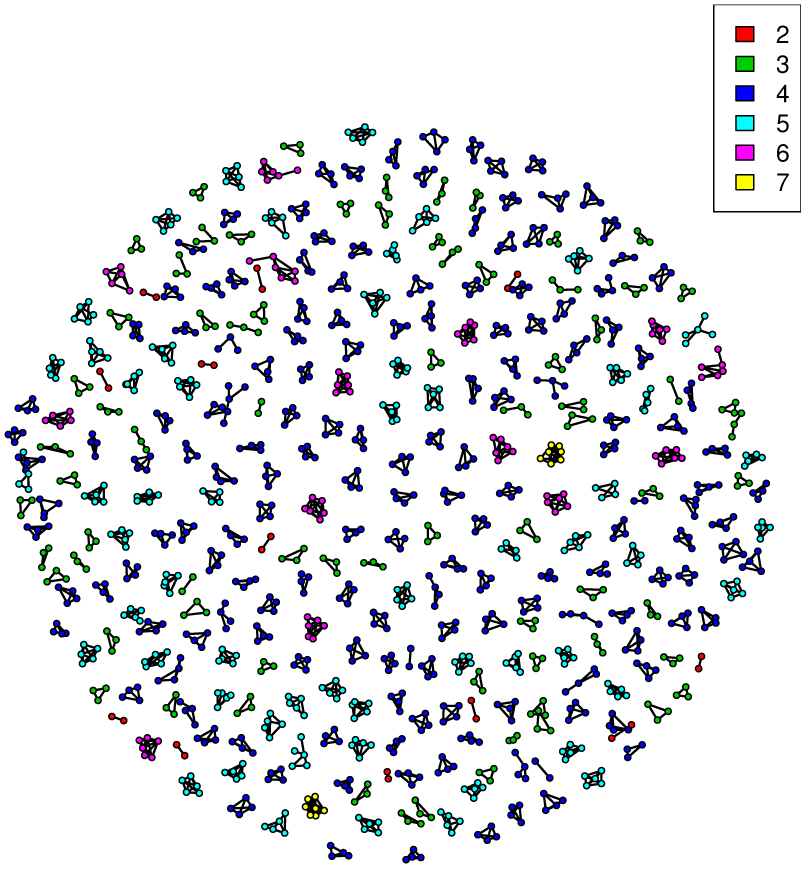
Observed within-household physical contact networks by household size (see color legend). Nodes represent household members and edges represent physical contacts.

Age, gender and household size were used to assign the role of child, mother and father to each household member. Two households were excluded from further analysis due to the assigning issue of a grandparent and a same-sex couple. The final data set thus consists of 316 households including 1259 participants.

Table 1 summarizes the proportion of complete (i.e. fully connected) networks and the mean network density for the within-household physical contact networks by household size, distinguishing week from weekend days and regular from holiday periods. The network density is defined as the ratio of the number of observed edges to the number of potential edges.

**Table 1.** Proportion of complete networks and mean network density, stratified by household size, for the observed within-household physical contact networks, comparing week and weekend days (top) and regular and holiday periods (bottom).

| HH size | Week |  |  | Weekend |  |  |
| --- | --- | --- | --- | --- | --- | --- |
|  | Nr. HHs | Proportion complete | Mean density | Nr. HHs | Proportion complete | Mean density |
| 2 | 9 | 1.00 | 1.00 | 3 | 1.00 | 1.00 |
| 3 | 53 | 0.91 | 0.96 | 19 | 0.74 | 0.88 |
| 4 | 111 | 0.77 | 0.93 | 48 | 0.85 | 0.96 |
| 5 | 39 | 0.64 | 0.90 | 18 | 0.78 | 0.95 |
| $\geq 6$ | 13 | 0.46 | 0.85 | 3 | 1.00 | 1.00 |
| Total | 225 | 0.77 | 0.93 | 91 | 0.82 | 0.94 |
| HH size | Regular period |  |  | Holiday period |  |  |
|  | Nr. HHs | Proportion complete | Mean density | Nr. HHs | Proportion complete | Mean density |
| 2 | 9 | 1.00 | 1.00 | 3 | 1.00 | 1.00 |
| 3 | 42 | 0.86 | 0.94 | 30 | 0.87 | 0.93 |
| 4 | 105 | 0.82 | 0.94 | 54 | 0.76 | 0.93 |
| 5 | 38 | 0.66 | 0.91 | 19 | 0.74 | 0.92 |
| $\geq 6$ | 12 | 0.50 | 0.84 | 4 | 0.75 | 0.98 |
| Total | 206 | 0.79 | 0.93 | 110 | 0.79 | 0.93 |

Overall, the type of day does not have a large impact on the contacts within households, however, the data suggest some decreasing connectedness with increasing household size, mainly on weekdays and during regular periods. For households of size 4, the observed proportion of complete networks is 0.77 on weekdays and 0.85 on weekend days.

Various measures of within-household clustering are defined in the SI text and Table S1 shows the high degree of physical contact clustering observed within households.

### Modeling within-household physical contact networks

We use ERGMs [19] to model the within-household physical contact networks. We explore the effect of relationships i.e. mixing between siblings, among children and their parents and between partners, gender-preferential mixing and age effects in children, and the effect of household size, distinguishing small (≤ 3 members), medium (4 members) and large (≥ 5 members) households (Table S2). We also explore the presence of higher-order dependency effects between members of the same household, such as clustering (see Table S1), by including in the model the number of isolate individuals, 2-stars, triangles and triangles in households of size ≥ 6. A 2-star is a person connected to two other household members and a triangle is a set of three household members such that all three are connected to each other.

The within-household physical contact networks were modeled separately for weekdays and weekends and the final ERGMs are presented in Table 2. The estimates shown in this table are log odds ratios and, hence, need to be exponentiated to obtain odds ratios. Note that the edge effect is estimated negative to counterbalance the large within-household edge effect, which is needed because our data does not include between-household contacts. For both types of days, the effects of gender-preferential mixing and the number of isolates were found to be non-significant (likelihood ratio test *p* = 0.5766 for weekdays). For weekend days, no significant effect of household size was found and the model was further reduced to an 8-parameter model (likelihood ratio test *p* = 0.5134). On weekdays, the odds of a physical contact occurring in a household of size ≤ 3 and ≥ 5 are estimated to be 2.10 and 0.67 times the odds of a physical contact occurring in a household of size 4, respectively. Thus, the network density for physical contacts decreases with increasing household size. Further, on both type of days, the odds of a physical contact between father and child is smaller than for any other pair except for older siblings, as the probability for siblings to make physical contact decreases with increasing age (Figure S3). For households of size ≤ 5, the odds of a physical contact that will complete a triangle is estimated to be 7.85 and 35.87 times the odds of a physical contact that will not complete a triangle on week and weekend days, respectively. This demonstrates the overall high degree of contact clustering within households. On weekdays, the degree of clustering is slightly lower in households of size ≥ 6 (conditional odds of 5.93).

**Table 2.**
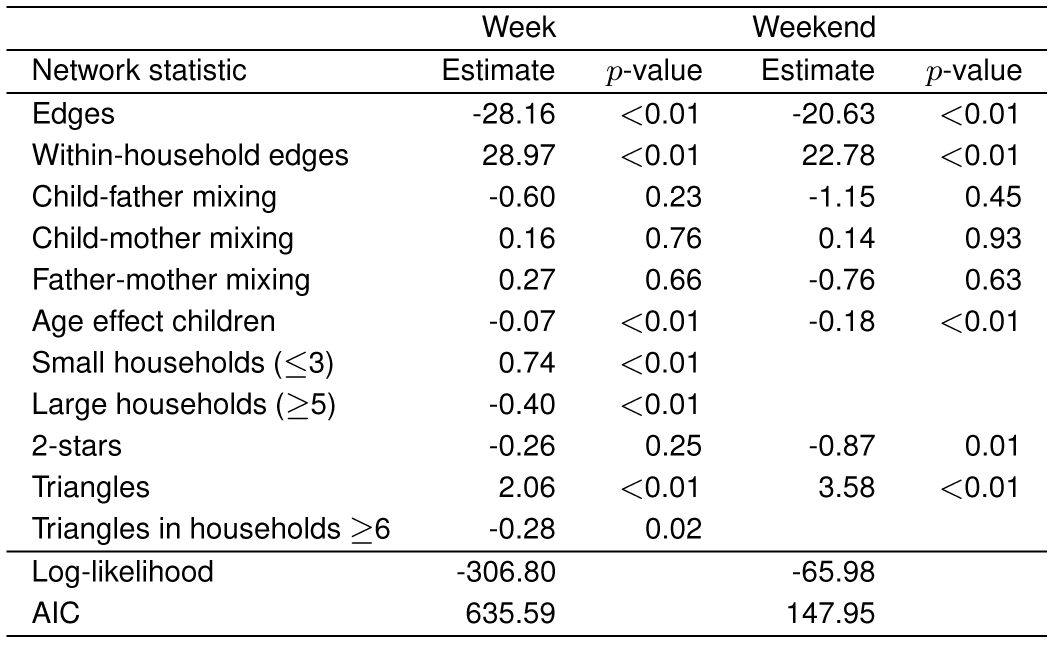
**ERGM for within-household physical contact networks on week- and weekend days: parameter estimates and Wald test ***p***- values, log-likelihood and AIC.**

Goodness-of-fit of the models is assessed by simulating new sets of physical contact networks from the fitted ERGM and by comparing specific contact network characteristics that are not included in the model, to the observed ones. We compare the proportion of complete networks, the mean network density and the proportion of observed versus potential triangles, by household size. Overall, the final ERGMs fit the data well as shown in Tables S3-S6 and Figures S4-S6.

### Epidemic spread in a community of households

We simulate the spread of a newly emerging infection in a closed fully susceptible population of households using a discrete-time chain binomial SIR (Susceptible - Infected - Recovered) model [27]. The 225 households from the contact survey that were analyzed using the weekday ERGM, are used to construct the community of households. We assume two levels of mixing similar to the households model in [7]: high-intensity mixing within households and low-intensity ‘background’ random mixing in the community i.e. between households. Two different configurations for within-household mixing are compared: random mixing and empirical-based mixing, where the latter refers to physical contact networks simulated from the fitted ERGMs. For each epidemic simulation, two sets of within-household contact networks are drawn from the ERGMs, one for a weekday and one for a weekend day, and those are kept fixed during the whole simulation.

Since we aim to study the effect of contact heterogeneity, we assume that susceptibility and infectiousness are invariant with age. Further, we assume that there is no latent period i.e. individuals are infectious immediately upon acquiring infection. At each time step, infected individuals recover with a constant probability of 0.22 such that the mean infectious period is approximately 3.5 days. Values for the transmission parameters are chosen in line with literature estimates for influenza based on household final size and symptom onset data (Table S7). The first day of the epidemic is randomly determined to be a week - or weekend day and is started by infecting three random individuals. The epidemic is then tracked until all infected individuals are recovered and no new infections have occurred.

### Scenario 1

Results from 1000 stochastic epidemic simulations are shown in Figures 2 and 3, S7 and S8. In these figures, small outbreaks defined as outbreaks with a final size of < 100 individuals that took less than 60 days, are excluded from display. The proportion of small outbreaks is significantly smaller in the random mixing setting compared to empirical-based mixing, 0.43 and 0.50, respectively (Fisher’s exact test, p-value: 0.0027).

**Fig 2.**
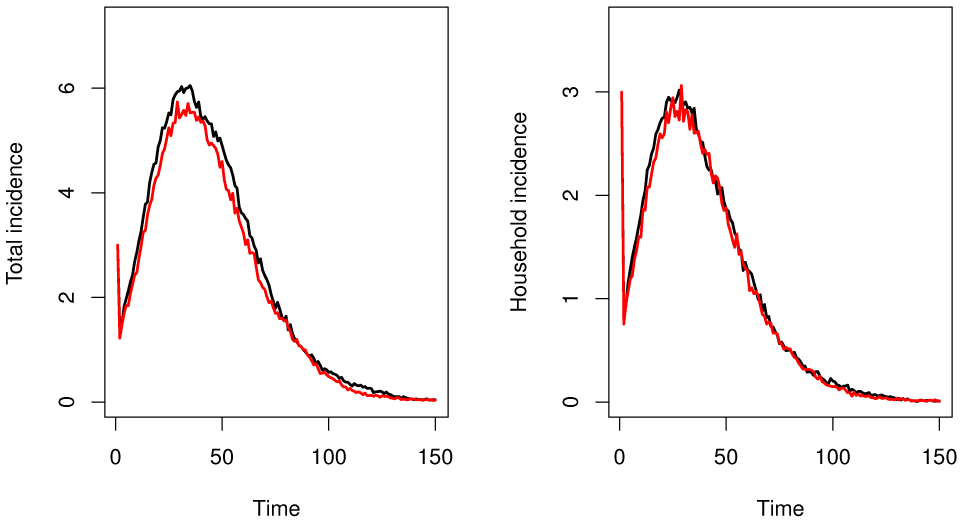
Mean infection incidence over time at the individual (left; number of newly infected individuals over time) and household level (right; number of newly infected households over time) assuming random (black) and empirical-based mixing (red) within households. Small outbreaks are excluded from display.

The mean proportion of individuals ultimately infected and the mean proportion of households infected is slightly larger under random mixing compared to realistic mixing: 0.39 [0.12, 0.56] vs. 0.36 [0.12, 0.53], and 0.70 [0.28, 0.88] vs. 0.67 [0.29, 0.86], respectively (Figure S8). Furthermore, the household attack rate, defined as the mean proportion of individuals infected per household [4] increases with household size under both settings (Figure 3).

**Fig 3.**
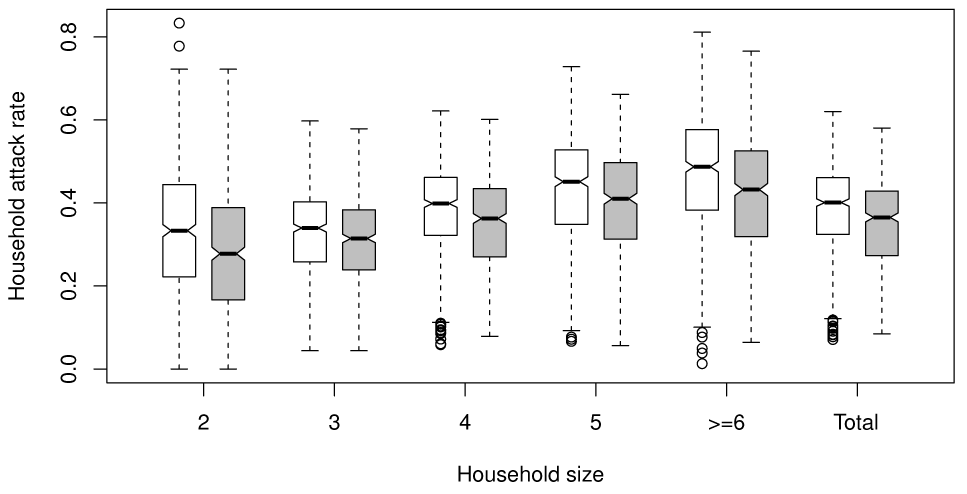
Household attack rates (mean proportion of infected individuals per household) by household size assuming random (white) and empirical-based mixing (gray) within households. Small outbreaks are excluded from display.

### Scenario 2

In the previous simulations, the small differences between the network model and the random mixing scenario could be due simply to different densities rather than to any particular characteristic of the network structure. In this setting we calibrate in order to make a more fair comparison between the two scenarios (see SI text). Figures 4-5, S9 and S10 present the results from 1000 simulations. Figure 4 shows the same epidemic dynamics over time and Figure 5 shows that the relation between household attack rate and household size is the same in both settings. Furthermore, there are barely any differences in mean final fraction of individuals (0.37 [0.13, 0.52] vs. 0.36 [0.12, 0.53]) and mean final fraction of households (0.68 [0.31, 0.86] vs. 0.67 [0.29, 0.86]; Figure S10). The proportion of small outbreaks is similar in both settings, 0.48 and 0.50, respectively (Fisher’s exact test, p-value: 0.3954).

**Fig 4.**
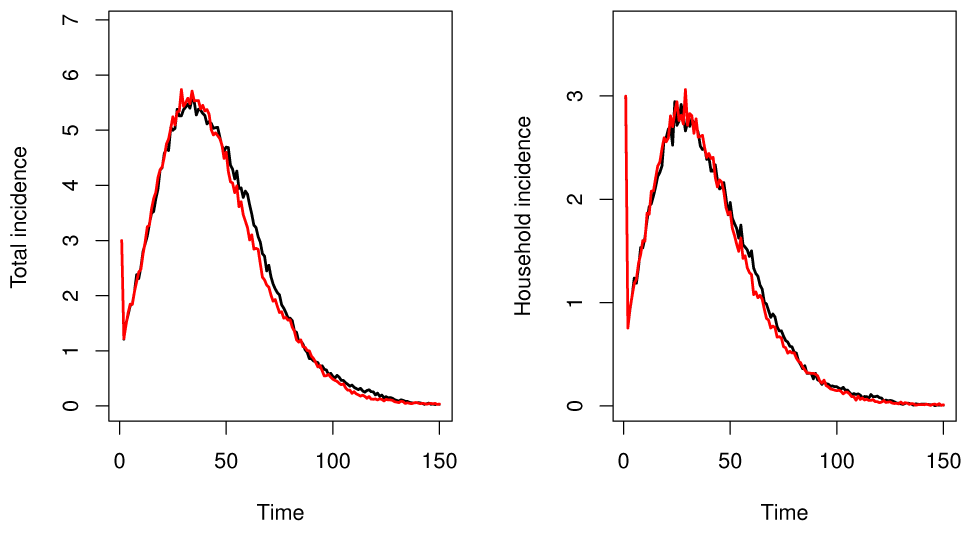
Mean infection incidence over time at the individual (left; number of newly infected individuals over time) and household level (right; number of newly infected households over time) assuming random (black) and empirical-based mixing (red) within households including a density scaling factor. Small outbreaks are excluded from display.

**Fig 5.**
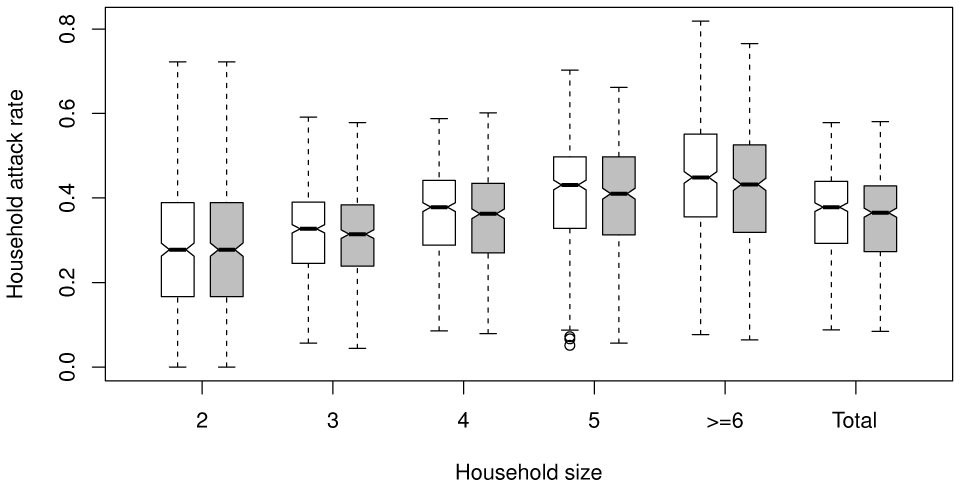
Household attack rates (mean proportion of infected individuals per household) by household size assuming random (white) and empirical-based mixing (gray) within households including a density scaling factor. Small outbreaks are excluded from display.

A more ‘extreme’ setting, focusing on physical contacts with a duration of more than 4 hours and assuming a higher within-household transmission rate, yields a lower incidence for empirical-based mixing regardless of correcting for the within-household density (see Figures S11-S14).

## Discussion

In this paper, we presented the first social contact study focusing specifically on contact networks within households. Inference of within-household contact networks in previous studies was based on egocentric contact data from the POLYMOD study [18] [14, 15] or on limited data in a very specific setting (rural Peru; [28]). Estimates of the proportion of complete networks inferred by Potter et al. [14, 15] ranged from 0.34 to 0.65 for households of size 4, and are thus smaller than the proportions that we observed (0.77 on weekdays and 0.85 on weekend days). The former estimates were based on partially observed within-household contact networks and, therefore, likely underestimated the true proportion of complete networks. For the purpose of studying household contacts, the current household-based survey design is considered an improvement upon the individual-based survey design (POLY-MOD study, [18]). We analyzed the household network data using ERGMs to assess the effect of factors such as role in the household, gender, children’s age and household size on physical contacts within households. We found that contacts between father and children are less likely than contacts between father and mother, mother and children and between siblings (except older siblings). This is in line with conclusions obtained by de Greeff et al. [29]. They analyzed data for pertussis in households with young infants and found that fathers were less likely to contract pertussis than other household members. Targeted vaccination of mothers and siblings was found to be most effective, as siblings are more likely to introduce an infection in the household. Further, we found that the contact density decreases and that the mean number of contacts increases with increasing household size (see Table 1 and S1). This implies that the mean contact degree is proportional to *z^w^* where *z* is household size and 0 *< w <* 1. This result supplements findings from studies on household epidemic data of close-contact infections [3, 29, 30] from a social contact data perspective. Finally, by simulating epidemics in a two-level SIR setting using literature-based influenza parameters, we found that solely incorporating contact heterogeneity has no impact on epidemic spread. This indicates that in this setting the assumption of random mixing between household members may be an adequate approximation of social contact behaviour for infections transmitted via close contacts. However, the results do suggest that accounting for the within-household contact density is of importance. Furthermore, we found that in a more extreme setting with intenser contacts and a higher within-household transmission rate a density correction is not sufficient to bridge the differences between both mixing assumptions. This suggests that informing mixing between household members with social contact data could impact modelling efforts in certain settings.

Our study has some limitations and assumptions. We assume that a contact occurred if it was reported by at least one household member. Thus, contacts forgotten by both members could result in an underestimation of the network density. Potter et al. [31] developed a model to deal with the issue of reporting error on network edges. However, given that the high reciprocity rates (98%) indicate a very good reporting quality of the survey, we believe such an adjustment will not have a large impact on our conclusions. Further, our results depend on the contact definition used to determine the within-household network links and cannot be generalized to the spread of any infectious disease. Based on the exploration of various contact definitions when using POLYMOD contact data to estimate age-specific varicella transmission rates [25], we opted to use physical contacts in this study as a surrogate of potential transmission events for close-contact infections such as influenza and smallpox. Though, even for two airborne infections, different networks may be appropriate because differing levels of interaction will be required to constitute an effective contact [32].

The methods in this paper could be extended in a number of ways which would be interesting topics of future research. We observed a relationship-specific heterogeneity in duration of contact (presented in Figure S2) and an impact of this duration on epidemic spread, which might be relevant for some diseases. The ERGM framework can be adapted to model ‘valued within-household contact networks’ [33], with the value of a contact determined by its total duration, and by weighting the transmission rates in the epidemic simulation model accordingly. It is also of potential interest to capture temporal dynamics of within-household contacts and to simulate the impact of contact formation and dissolution on the spread of infection [34, 35]. Combining time-use data with social contact data would allow to infer the potential timing of contacts with household members, and to estimate dynamic within-household contact networks. This would also be valuable to inform large-scale individual-based simulation models of infectious disease spread. Further, exploration of potential differences in the distribution of the generation interval in a random mixing setting versus empirical-based mixing is the topic of current research. Finally, combining the model for within-household contact networks developed in this paper with epidemic data from a similar community of households, would allow to improve estimates of age-specific heterogeneity in susceptibility and infectiousness for infections such as influenza [6].

This study provides unique insights into within-household contacts, considered to be important drivers of many close-contact infections. It is the first empirical evidence resulting from a large household contact survey supporting the use of the random mixing assumption in epidemic models incorporating household structure.

## Materials and Methods

### Exponential random graph model

Let ***Y*** denote the random adjacency matrix of an undirected network, where *Y*_*i,j*_ = *Y*_*j,i*_ = 1 if person *i* and *j* made physical contact and zero if not, and let Ω denote the support of **Y** i.e. the set of all obtainable networks. In an ERGM, the probability of observing a set of network edges is defined as follows:

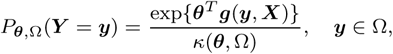

where ***g***(***y***,***X***) is a vector of network statistics that may depend on additional covariate information ***X***, with *θ* the corresponding vector of coefficients, and *κ*(*θ,Ω*) a normalizing factor. By using an alternative model specification (see SI text), *θ* can be interpreted as the increase in the conditional log-odds of the network, per unit increase in the corresponding component of ***g***(**y**,***X***), resulting from switching a particular dyad *Y*_*i,j*_ from 0 to 1 while leaving the rest of the network fixed at 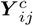.

We infer on the processes driving physical contacts between household members by incorporating network statistics based on nodal covariate information (see Table S2). Although our analysis is focused on within-household contact networks, we fit a single ERGM including all households. We include in our model, described in detail in the SI text, a household effect which captures the tendency to contact others in one’s own household. Because there are no between-household contact reports present in our survey, the coefficient for the household preference effect should be estimated to be extremely large, so the probability of between-household contact is essentially zero.

Approximate maximum likelihood estimates are obtained using a stochastic Markov Chain Monte Carlo (MCMC) algorithm [36]. MCMC estimation is performed with the "ergm" package in R [37, 38] that is part of the "statnet" suite of packages for statistical network analysis [39–41].

## Epidemic simulations

The discrete-time chain binomial SIR model is defined in more detail in the SI text.

## ACKNOWLEDGMENTS

This work benefited from discussions held with Ken Eames, Lorenzo Pellis, Chiara Poletto and James Wood. This project has received funding from the European Research Council (ERC) under the European Union’s Horizon 2020 research and innovation programme (grant agreement 682540 - TransMID). Support from the IAP Research Network P7/06 of the Belgian State (Belgian Science Policy) is gratefully acknowledged. NG was supported by a postdoctoral grant from the AXA Research Fund. NH gratefully acknowledges support from the University of Antwerp scientific chair in Evidence-Based Vaccinology, financed in 2009–2016 by a gift from Pfizer and GSK. ES acknowledges support from a Methusalem research grant from the Flemish government. LW is supported by the Research Foundation Flanders (FWO, project G043815N). AT acknowledges support from BOF funding from the University of Antwerp. This research was supported by the Antwerp Study Centre for Infectious Diseases (AS-CID).

